# Human airway lineages derived from pluripotent stem cells reveal the epithelial responses to SARS-CoV-2 infection

**DOI:** 10.1101/2021.07.06.451340

**Authors:** Ruobing Wang, Adam J. Hume, Mary Lou Beermann, Chantelle Simone-Roach, Jonathan Lindstrom-Vautrin, Jake Le Suer, Jessie Huang, Judith Olejnik, Carlos Villacorta-Martin, Esther Bullitt, Anne Hinds, Mahboobe Ghaedi, Rhiannon B. Werder, Kristine M. Abo, Andrew A. Wilson, Elke Mühlberger, Darrell N. Kotton, Finn J. Hawkins

## Abstract

There is an urgent need to understand how SARS-CoV-2 infects the airway epithelium and in a subset of individuals leads to severe illness or death. Induced pluripotent stem cells (iPSCs) provide a near limitless supply of human cells that can be differentiated into cell types of interest, including airway epithelium, for disease modeling. We present a human iPSC-derived airway epithelial platform, composed of the major airway epithelial cell types, that is permissive to SARS-CoV-2 infection. Subsets of iPSC-airway cells express the SARS-CoV-2 entry factors *ACE2* and *TMPRSS2.* Multiciliated cells are the primary initial target of SARS-CoV-2 infection. Upon infection with SARS-CoV-2, iPSC-airway cells generate robust interferon and inflammatory responses and treatment with remdesivir or camostat methylate causes a decrease in viral propagation and entry, respectively. In conclusion, iPSC-derived airway cells provide a physiologically relevant *in vitro* model system to interrogate the pathogenesis of, and develop treatment strategies for, COVID-19 pneumonia.

**Highlights and eTOC blurb:**

- Subsets of human iPSC-airway epithelial cells express SARS-Co-V entry factors *ACE2* and *TMPRSS2*.
- iPSC-airway cells are permissive to SARS-CoV-2 infection via multiciliated cells.
- SARS-CoV-2 infection of iPSC-airway leads to a robust interferon and inflammatory response.
- iPSC-airway is a physiologically relevant model to study SARS-CoV-2 infection.

## INTRODUCTION

Infection with the severe acute respiratory syndrome coronavirus 2 (SARS-CoV-2), a zoonotic positive sense RNA virus, has been a major source of worldwide morbidity and mortality since its emergence, with current analyses indicating that it was the third leading cause of death in the United States in 2020^1^. The major morbidity and mortality from SARS-CoV-2 results from infection of the respiratory system causing COVID-19 pneumonia. While respiratory failure and Acute Respiratory Distress Syndrome (ARDS) develop as consequences of infection involving the gas-exchanging alveolar compartment of the lung, the initial infection involves the nasal and subsequently airway epithelium^2, 3^. Viral entry into cells is achieved by binding of the SARS-CoV-2 spike (S) glycoprotein to the human angiotensin-converting enzyme (ACE2) receptor and subsequent processing of the S protein by proteases, including transmembrane protease, serine 2 (TMPRSS2)^4, 5^. *ACE2* expression levels decrease along the respiratory tract with the highest expression in the nose and lowest expression in the distal lung^3^. This gradient suggests a paradigm of initial infection in the susceptible nasal epithelium with subsequent propagation to the ACE2-expressing cells of the airways and alveoli^3^. While it is clear that the airway epithelium is a major target of SARS-CoV-2^3, 5–7^, the mechanisms by which the airway responds to infection, triggers an immune response, and leads to a wide variation of disease severity is unclear and of major interest. A comprehensive understanding of the pathogenesis of SARS-CoV-2 in the airways is necessary to advance prognostic tools and therapeutics to combat COVID-19.

To enhance our understanding of SARS-CoV-2 infection in the airways, *in vitro* models that are permissive to SARS-CoV-2 infection and recapitulate the pathology of COVID-19 respiratory tract infection are required^8^. SARS-CoV-2 can replicate in cell lines such as Calu-3^6^ and Caco-2 cells^7^. However, these cancer cell lines have lost much of their tissue-specific cell programs^9^. Vero and Vero E6 cells, immortalized kidney cell lines derived from an African green monkey, are widely used in SARS-CoV-2 research, since they are highly infectible by SARS-CoV-2^10^, amenable to high-throughput approaches, and were used in the original studies that identified ACE2 as the receptor for SARS-CoV-1^11^. However, the responses of these cell lines may not be representative of human or lung epithelial responses to infection^9^. Human primary bronchial epithelial cells (HBECs) can be differentiated in well-established protocols into a mucociliary epithelium in an air-liquid interface (ALI) culture format that is considered the gold standard *in vitro* model of human airway epithelial biology^12^. Multiple studies have demonstrated the permissiveness of primary HBECs to SARS-CoV-2 infection and documented an epithelial immune response^12–14^. Relevant to future SARS-CoV-2 studies, there are aspects of HBECs that are limiting including: 1) limited access to human tissues from diverse populations/diseases, 2) finite *in vitro* life-span and differentiation capacity over time, and 3) limited success with gene-editing approaches to interrogate the role of specific genes or variants of interest^15, 16^. Therefore, access to an additional physiologically relevant, renewable source of human airway epithelial cells that are permissive to SARS-CoV-2 infection has the potential to facilitate mechanistic studies into SARS-CoV-2 pathogenesis and drug development strategies.

Induced pluripotent stem cells (iPSCs) have several features relevant to COVID-19 disease modeling including: 1) the near-limitless supply of cells, 2) their retention of the genetic profile of the individual from which they were generated, 3) the capacity to differentiate under appropriate conditions into tissue-specific cell types relevant to SARS-CoV-2 infection, including both proximal and distal lung epithelia, and 4) amenability to gene-editing^17, 18 19 20^. Through the *in vitro* recapitulation of developmental milestones via directed differentiation, we and others have generated iPSC-derived airway and alveolar epithelial cells^21–37^. We recently demonstrated the feasibility of using iPSC-derived alveolar epithelial type 2 cells (iAT2s) to model SARS-CoV-2 infection^9, 38 39^. iAT2s were permissive to infection with SARS-CoV-2, which induced a rapid inflammatory response characterized by secretion of NFκB-induced cytokines and moderate, delayed type I and III interferon responses. Treatment of infected iAT2s with remdesivir, an inhibitor of the SARS-CoV-2 RNA-dependent RNA polymerase, resulted in a decrease in viral replication, demonstrating the potential of the iPSC platform for drug testing^38^. In terms of the initial target of SARS-CoV-2 infection, we recently developed methods to direct the differentiation of human iPSCs into airway basal cells (iBCs)^40^. Basal cells are the major stem cells of the airway epithelium^41^; and iBCs are molecularly similar to their endogenous counterparts and express canonical basal cell markers including the transcription factors TP63 and NKX2-1 and the surface receptor, NGFR^40^. iBCs are functionally similar to primary basal cells based on their multi-lineage differentiation capacity in ALI cultures, forming pseudostratified epithelia composed of basal, secretory, and multiciliated cells. These iPSC-derived airway ALI cultures (iPSC-airway) are similar in morphology and composition to primary HBEC-derived ALI cultures^40^.

Here we report the successful infection of iPSC-airway by SARS-CoV-2 and characterization of the resulting epithelial response. We demonstrate that multiple cell types in our iPSC-airway model express the viral entry factors *ACE2* and *TMPRSS2*, and we find that multiciliated cells are the initial primary targets of infection. Following infection, we identified robust type I and type III interferon responses that contrast with our findings in SARS-CoV-2 infection of iAT2 cells^38^. In addition, we observed an inflammatory response implicating signaling via the NFκB pathway. Finally, treatment with remdesivir and camostat methylate caused a decrease in virus in our platform, demonstrating the feasibility of the platform for drug screening assays.

## RESULTS

### Human iPSC-airway expresses SARS-CoV-2 entry factors

We previously generated from a healthy donor a bifluorescent reporter iPSC line that carries a tdTomato coding sequence targeted to the *TP63* locus and a GFP sequence targeted to the *NKX2-1* locus^21, 40^. When differentiated using a previously published 6 stage protocol **(Figure 1A)**^40^, this BU3 NKX2-1^GFP^;P63^tdTomato^ dual reporter line (hereafter BU3 NGPT) allows the sequential identification, and purification via flow cytometry, of lung progenitors (NKX2-1^GFP+^), airway progenitors (NKX2-1^GFP+^/TP63^tdTomato+^), and finally basal cells (NKX2-1^GFP+^/TP63^tdTomato+^/NGFR^+^)(**Figures S1A-C**). As an alternative to the reporter-based approach we also developed a surface marker-based strategy in which NKX2-1+ lung progenitors are identified using CD47^hi^/CD26^neg^ sorting^21^ followed by NGFR+ iBC purification. Following these established differentiation protocols, we differentiated BU3 NGPT and a previously described human iPSC line (hereafter “1566”) using reporter-based sorting or surface marker-based sorting, respectively **(Figures S1D-F)**^40^. Plating of purified iBCs from both iPSC lineages to a Transwell ALI culture format (removing media from the apical chamber and exposing the cells to air) resulted in their differentiation to a mucociliary airway epithelium (**Figures 1B-C**).

**Figure 1.**
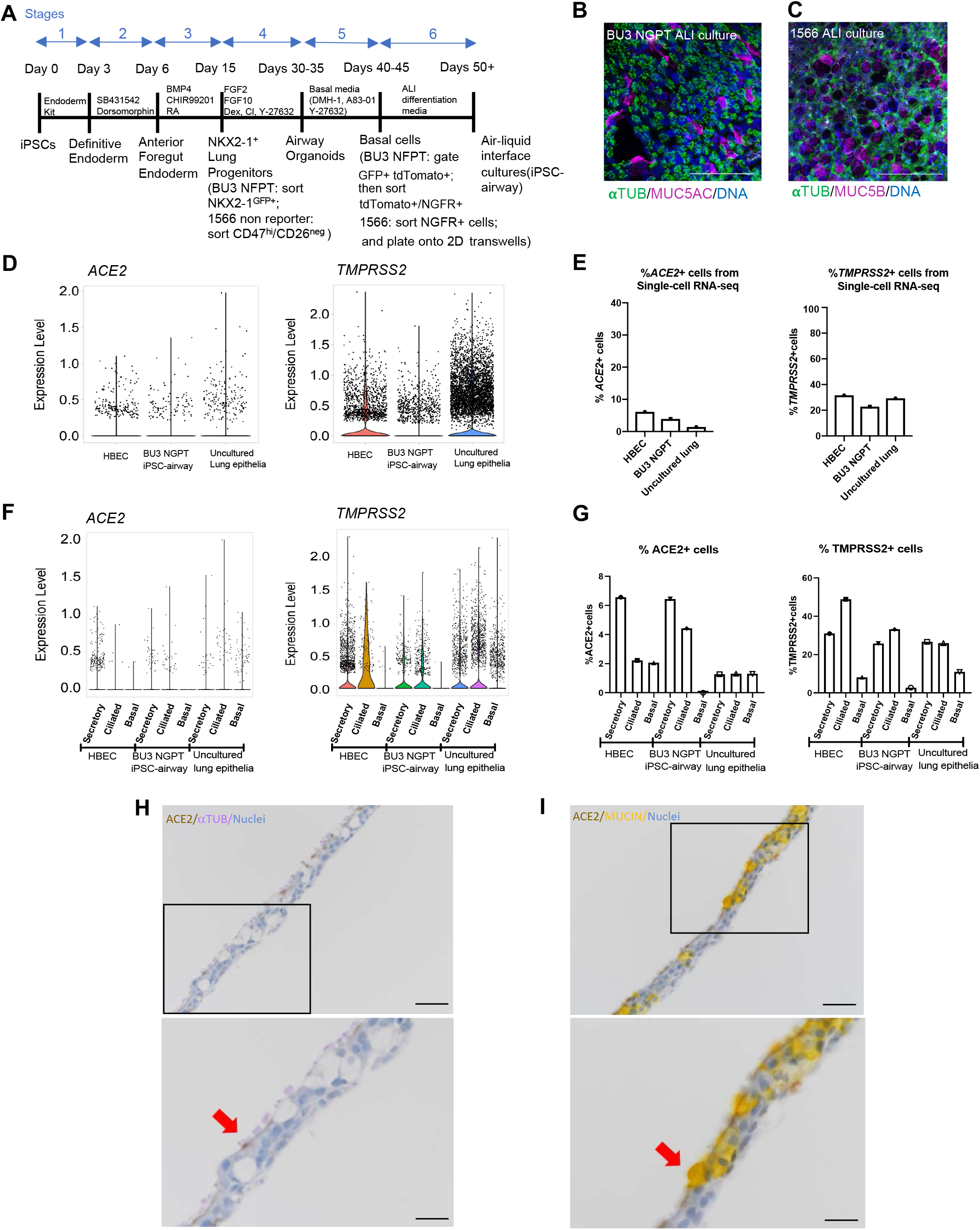
iPSC-derived airway cells express SARS-CoV-2 entry factors *ACE2* and *TMPRSS2*. A) Schematic of the 6-stage differentiation protocol of generating iPSC-airway. B) Immunofluorescence analysis of BU3 NGPT iPSC-airway stained with anti-α-TUBULIN and MUC5AC (scale bar = 100 μm). Nuclei are stained with HOESCHT (blue). C) Immunofluorescence analysis of 1566 iPSC airway, stained with anti-α-TUBULIN and MUC5B (scale bar = 100μm). Nuclei are stained with DAPI (blue). D-G) scRNA-seq analysis of HBEC^40^, iPSC-airway (BU3 NGPT)^40^, and freshly isolated uncultured lung epithelia^42^. D) Violin plots of *ACE2* and *TMPRSS2* expression. E) The percentage of *ACE2* and *TMPRSS2* positive cells in each dataset^40^. F) Violin plots of *ACE2* and *TMPRSS2* expression by cellular type in each dataset. G) Comparison of the percentage of *ACE2* and *TMPRSS2* positive secretory, multiciliated, and basal cells in each dataset. H-I) Immunohistochemistry staining showing the localization of ACE2 (DAB), α-TUBULIN (purple, left panels) MUCIN (yellow, right panels) in iPSC-derived airway (BU3 NGPT) counterstained with hematoxylin (20x, scale bar =50μm). Lower panels are zoomed-in images of the black box in the upper panels. The red arrows indicated cells co-expressing ACE2/ α-TUBULIN (left) and ACE2/MUCIN (right) (scale bar =25μm).

To interrogate the feasibility of applying the iPSC-airway system to model SARS-CoV-2 infection of the airway, we first examined the expression levels of SARS-CoV-2 entry factors, *ACE2* and *TMPRSS2*. To do so, we compared previously published single-cell RNA-sequencing (scRNA-seq) datasets: airway epithelium derived from BU3 NGPT iBCs ALI cultures (BU3 NGPT iPSC-airway)^40^, and primary HBECs differentiated in ALI cultures from a healthy donor^40^. We next integrated these datasets with a recently-published scRNA-seq dataset of freshly isolated (uncultured) lung epithelial tissue from 6 donors without underlying lung disease^42^. Analyzing all airway epithelia in each of these samples, we identified subsets of cells expressing *ACE2* and *TMPRSS2* in BU3 NGPT iPSC-airway, HBEC-derived ALI cultures (hereafter HBEC), and uncultured lung epithelia **(Figure 1D)**. We quantified the percentage of cells expressing *ACE2* and *TMPRSS2* transcripts: *ACE2* was detected in 6.1% of HBEC, 3.9% of BU3 NGPT iPSC-airway, and 1.4% of uncultured lung epithelia; *TMPRSS2* was present in 31.6% of HBEC and 29.3% of uncultured lung epithelia, compared to 22.8% of BU3 NGPT iPSC-airway (**Figure 1E**).

Next, we examined the cell-type specific expression of *ACE2* and *TMPRSS2* across these datasets **(Figure 1F)**. The annotated clusters and expression of canonical markers used to define secretory, multiciliated, and basal cells are shown in **Figure S1G**. The percentage of *ACE2*+ secretory cells were 6.6%, 6.5%, and 1.3% in HBEC, BU3 iPSC-airway, and uncultured lung epithelia, respectively **(Figure 1G)**. The percentage of *ACE2*+ multiciliated cells were 2.2%, 4.4%, and 1.3%, and the percentage of *ACE2*+ basal cells were 2.1%, 0%, and 1.3% in HBEC, BU3 iPSC-airway, and uncultured lung epithelia, respectively **(Figure 1G)**. For *TMPRSS2*, 31.0%, 25.8%, and 26.7% of secretory cells, 48.9%, 33.2%, and 26.0% of multiciliated cells, and 8.2%, 2.8%, and 11.0% of basal cells from HBEC, iPSC-airway, and uncultured lung epithelia, respectively, were positive **(Figure 1G)**. Taken together, while there are apparent differences in *ACE2* and *TMPRSS2* expressions when comparing *in vitro* platforms to *in vivo*, and iPSC-airway to primary cells, in general *ACE2* is expressed in small subpopulations of cells across all three platforms. Furthermore, similar to a recent finding that cells that co-express *ACE2* and *TMPRSS2* are enriched in pathways related to viral infection and immune response^43^, we also found that the top enriched genes, ranked by Z-score, in iPSC-airway *ACE2*+ cells (vs *ACE2*-cells) included interferon stimulated genes such as *IFI27, IFIT1, RSAD2, ISG15, MX1 IFITM2, IFIT3* **(Table S1)**.

Finally, we examined ACE2 protein expression and localization in iPSC-airway cells (**Figures 1H-I**). In concordance with our scRNA-seq data, ACE2 was detected in subsets of α-TUBULIN+ multiciliated and MUC5AC+ secretory cells, and was apically localized (**Figures 1H-I**). Taken together, we demonstrate that SARS-CoV-2 entry factors *ACE2* and *TMPRSS2* are expressed in multiple lineages of iPSC-airway epithelial cells.

### iPSC-airway is permissive to SARS-CoV-2 infection

To determine whether iPSC-airway is permissive to SARS-CoV-2 infection, iPSC-airway cultures from BU3 NGPT and 1566 were differentiated following the airway differentiation protocol as described above. After approximately 21 days in ALI culture, the cells were infected from the apical surface with purified SARS-CoV-2 particles **(Figure 2A)**. Viral infection was detected by immunofluorescence analysis (IFA) using an antibody directed against the SARS-CoV nucleoprotein (N) with cross-reactivity against SARS-CoV-2 N protein (**Figures 2B-G)** and by RT-qPCR of the *N*-transcript (**Figures 2H, S2A-D**). We observed a dose-dependent increase in viral *N* transcript with increasing multiplicity of infection **(Figure S2A)**. MOI of 4 was selected for the subsequent studies. At 1 day post infection (dpi), an average of 6.87% (SEM 0.548%) of BU3 NGPT iPSC-airway cells were N + by IFA **(Figures 2B-C)**. In 1566 iPSC-airway infected with SARS-CoV-2, 11.21% (SEM 1.075) of cells were N + by IFA at 1 dpi **(Figures 2D-E)**. RT-qPCR of both BU3 NGPT and 1566 iPSC-airway confirmed the presence of *N-*transcript at 1 and 3 dpi (**Figures 2H, S2D)**. In 5 separate infections of BU3 NGPT iPSC-airway and 3 separate infections of 1566 iPSC-airway, we detected an average N transcript of at 1 dpi of 2.64×10^7^ (fold change over mock) and 2.44×10^7^ (fold change over mock), respectively (data not shown). We then followed the course of SARS-CoV-2 infection of iPSC-airway over time. In both iPSC lines, peak infection was detected a 1 dpi and decreased approximately 10-fold by 3 dpi (**Figures 2H**, **S2C-D**). Cellular toxicity was suggested at 3 dpi by lower cell density, patches devoid of nuclei, an increase in trypan-blue labeled non-viable cells and an increase in fragmented nuclei **(Figure 2G, I, S2E)**. To determine whether infectious virus was released from the infected iPSC-airway epithelium viral titers were performed on apical washes and basolateral medium from 1 and 3 dpi samples. Viral particles were mainly released form the apical surface (**Figure 2J**) which is in line with the transmission route of SARS-CoV-2 and similar to findings reported using primary HBECs^44^. We also observed increased shedding of infectious virus from both apical and basolateral compartments from 1 to 3 dpi (**Figure 2J**) while intracellular *N*-transcripts decreased.

**Figure 2.**
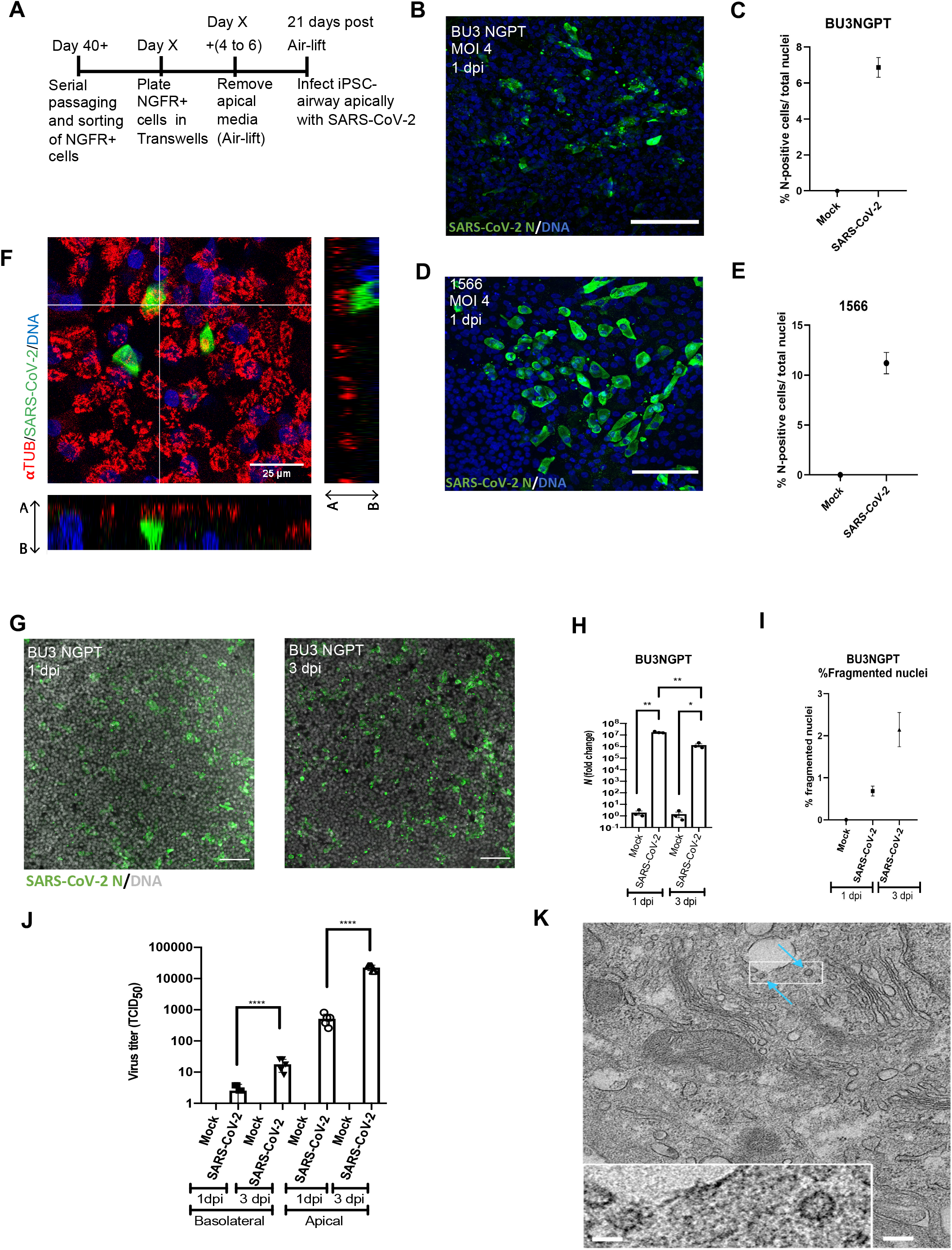
iPSC-derived airway is permissive to SARS-CoV-2 infection with time-dependent restriction in viral growth. A) Schematic of the protocol to iPSC-airway with SARS-CoV-2. B-E) Immunofluorescence and quantification of viral nucleoprotein+ (SARS-CoV-2 N, green) cells in BU3 NGPT (B-C) and 1566 (D-E) cells at 1 dpi (40x, scale bar=100μm). BU3 NGPT mean nucleoprotein+ cells =6.87%±0.548 (SEM). 1566 mean nucleoprotein+ cells = 11.21% ± 1.075 (SEM). F) Confocal immunofluorescence microscopy of BU3 NGPT with antibodies against SARS-CoV-2 N positive (green) cells and α-TUBULIN (red). Nuclei are stained with DAPI (blue). G) Immunofluorescence of infected iPSC airway (BU3 NGPT) at 1 and 3 dpi, labeled with anti-SARS-CoV-2 N antibody and nuclei labeled with DAPI (20x, scale bar =100um). H) RT-qPCR of viral N gene expression of iPSC-airway (BU3 NGPT) at 1 and 3 dpi (n = 3 Transwells per sample). Fold change expression compared to mock [2^−ddCt^] after 18S normalization is shown. I) Percent of fragmented nuclei detected at 1 and 3 dpi in iPSC-airway (BU3 NGPT)(1 dpi=mean 0.69±0.12% (SEM) and 3 dpi (mean 2.14±0.32 (SEM)). J) Viral titers from apical washes and basolateral media at 1 and 3 dpi from iPSC-airway (BU3 NGPT) compared to mock. K) Transmission electron micrograph of SARS-CoV-2 infected iPSC-airway (1566) at 1 dpi. Blue arrows indicate viral particles (Scale bar= 200nm, 50nm in enclosed box)

To study the cellular tropism of SARS-CoV-2 in our iPSC-airway system, we performed IFA with 1 dpi samples using antibodies directed against SARS-CoV nucleoprotein and canonical markers of multiciliated (acetylated tubulin, ACT), basal (TP63), or secretory cells (MUC5AC). By confocal microscopy, all cells that co-expressed a lineage marker and viral nucleoprotein were ACT+ multiciliated cells **(Figure 2F)**. We did not identify co-localization of viral nucleoprotein with MUC5AC+ secretory **(Figure S2F)** or TP63+ basal cells (not shown). The observation that multi-ciliated cells are the predominant initial airway target cells is in agreement with previous studies in SARS-CoV-2 infected primary airway cultures^3, 45^. Ultrastructural analysis of SARS-CoV-2 infected iPSC-airway by transmission electron microscopy (TEM) revealed the presence of intracellular viral particles, confirming productive viral infection **(Figure 2K**). Taken together, our findings indicate iPSC-airway cells from two different individuals are permissive to SARS-CoV-2 infection and multiciliated cells are the initial target cells. Though there may be increased virion release or potential extracellular accumulation on the apical surface of the infected cells at 3 dpi, the infection of the epithelium peaks prior to 3 dpi with declining intracellular viral presence over time, associated with cytotoxicity.

### SARS-CoV-2 infection stimulates an epithelial-intrinsic interferon and inflammatory response in iPSC-airway

Having demonstrated the permissiveness of the iPSC-airway system to SARS-CoV-2 infection, we next sought to assess the global transcriptomic response to infection. To do so, we performed bulk RNA-Sequencing (RNA-Seq) of SARS-CoV-2 infected iPSC-airway (BU3 NGPT) at 1 and 3 dpi, compared to time-matched mock-infected controls (n=3 replicates at each time point) **(Figure 3A)**. Principal component analysis (PCA) indicated that the predominant variance in global gene expression (PC 1; 42.6% variance) was explained by the infection state of the cells, with less variance (PC2; 20.2% variance) explained by time in culture **(Figure 3B, Table S2**. For example, 529 genes and 5134 genes were differentially expressed at 1dpi and 3 dpi, respectively, compared to time-matched, mock-infected samples (FDR) < 0.05). Comparing 3 to 1 dpi, 6136 genes were differentially expressed, with 2853 upregulated and 3283 downregulated genes **(Figure 3B, Table S2)**.

**Figure 3.**
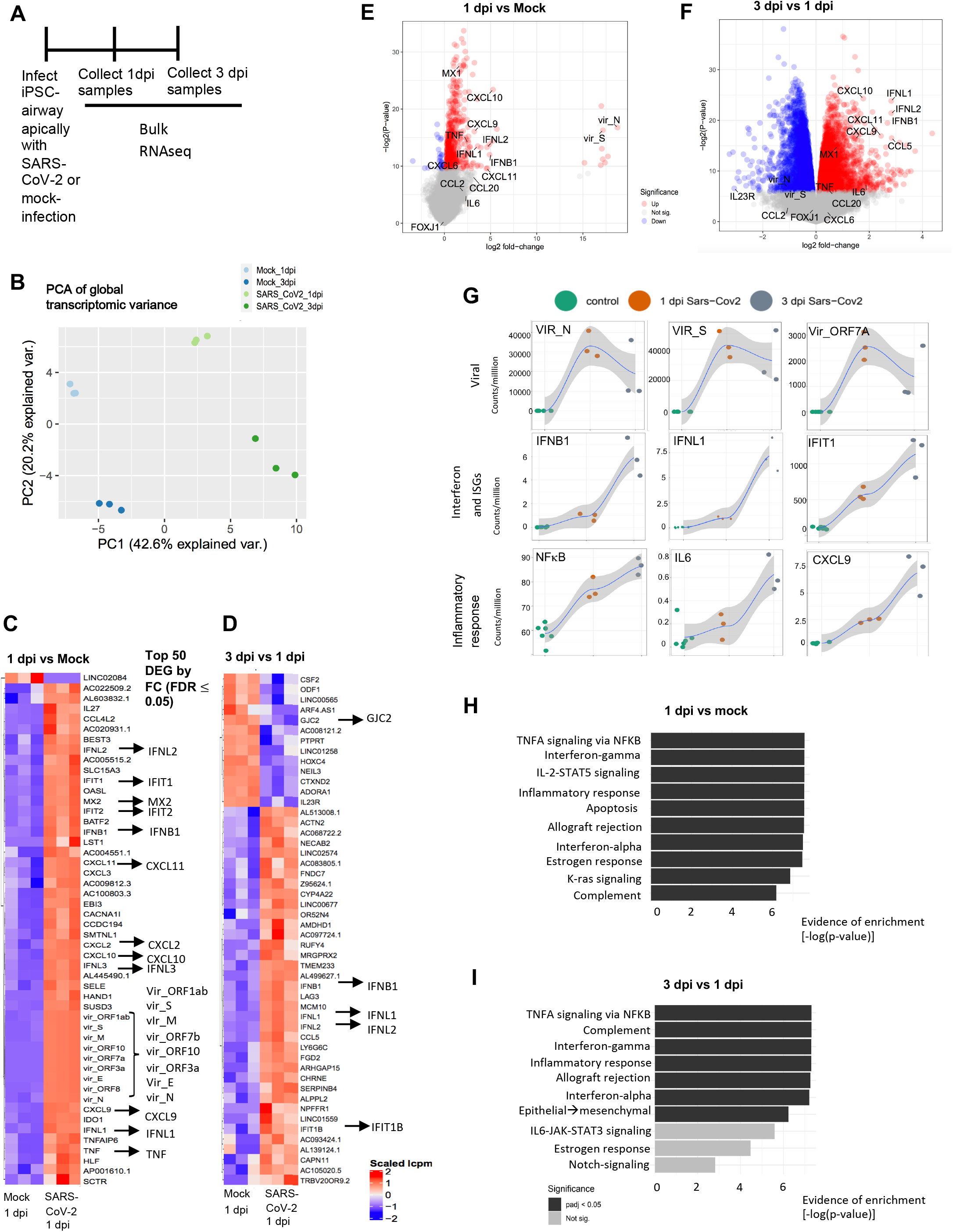
Transcriptomic analysis of SARS-CoV-2 infected iPSC-airway shows robust interferon response. A) Schematic of the experimental design to compare SARS-CoV-2 infected iPSC-airway samples (BU3 NGPT) at 1 and 3 dpi to mock controls by RNA-seq (n=3 Transwells). Figures 4B-I are analyses of this experiment. B) PCA comparing PC1 vs PC2 of the samples described in A. C) The top 50 DEGs ranked by fold change (FDR<0.05) of 1 dpi vs mock. D) The top 50 DEGs ranked by fold change (FDR<0.05) of 3 dpi vs 1 dpi. E) Volcano plots of differentially expressed genes in 1 dpi versus mock for iPSC-airway. F) Volcano plots of differentially expressed genes in 3 dpi versus 1 dpi for iPSC-airway. G) Local regression (LOESS) plots of viral, interferon and ISG, and inflammatory gene expression levels quantified by RNA-seq normalized expression (counts per million reads). H) Gene set enrichment analysis (GSEA) of the top ten upregulated gene sets in 1 dpi versus mock for iPSC-airway (black color indicates statistical significance; padj < 0.05). I) Gene set enrichment analysis (GSEA) of the top ten upregulated gene sets in 1 dpi versus 3 dpi iPSC-airway (black color indicates statistical significance; padj < 0.05)

Given the large number of genes changing after infection, we analyzed the most robust transcriptomic changes by focusing on the top 50 DEGs ranked by logFC (and filtered by FDR < 0.05) at 1 dpi (vs mock) (**Figure 3C**), 3 dpi (vs mock) (**Figure S3A**), and 3 dpi vs 1 dpi (**Figure 3D**). We found that 9/50 (18%) of the top DEGs at 1 dpi were viral transcripts as expected (logFC; p < 0.05) **(Figures 3C, 3E, S3A)**. At 1 dpi, genes associated with type I (including *IFNB1*), type III interferon (including *IFNL1, INFL3*), and downstream interferon-stimulated genes (ISGs) (*IFIT2, MX2, MX1*) were upregulated **(Figures 3C, E)**. Furthermore, genes encoding cytokines involved in the recruitment and differentiation of immune cells, including *CXCL9*, *CXCL10* and *CXCL11*, were also upregulated at 1 dpi compared to mock-infected samples **(Figures 3C, 3E)**. From 1 to 3 dpi, there was a decrease in viral gene expression, including transcripts from *N, M, E*, and *ORF3a*, consistent with RT-qPCR N transcript profiles (**Table S1**, FDR < 0.05). This was accompanied by further upregulation of type I and type III interferon genes (*IFNB*, *IFNL1, IFNL2*) and ISGs (*IFIT1B, MX1*), suggesting increasing interferon response over time **(Figures 3D, 3F)**. Furthermore, there was an upregulation of inflammatory mediators (*CXCL9, CXCL10, CXCL11)* and a modest increase in *IL6* expression from 1 to 3 dpi (**Figure 3F**).

The expression kinetics of viral transcripts, interferon genes, ISGs, and genes involved in the inflammatory response (including NFκB and IL-6) from 1 dpi to 3 dpi were also visualized using local regression (LOESS) plots (**Figure 3G, S3E)**. In addition to upregulation of type I/III interferons and ISGs as described above, there was also a time-dependent increase in viral sensors (*DDX58, TLR3*) and adaptor molecule (*MYD88*) **(Figure S3E)**. We also observed an increase in chemokines important for T and NK cell recruitment (*CXCL9, CXCL10, CXCL11*), *IL-23* implicated in the IL-17 pathway, *CXCL8* important for neutrophil migration, as well as *TNFAIP3* **(S3E)** <u>from 1 to 3 dpi</u>. CCL2, which is important for macrophage recruitment, was not upregulated **(S3E)**. Consistent with the findings that inflammation is a hallmark of SARS-CoV-2 infection^46–48^, gene set enrichment analysis (GSEA) based on the DEGs at 1 dpi (vs. mock) suggested enrichment of the following pathways; “TNFA signaling via NFκB”;, “IL-2-STAT5 signaling”; “Inflammatory response”; and “Complement” **(Figure 3H)**. Furthermore, GSEA also showed enrichment of “interferon-gamma” and “interferon-alpha” pathways, consistent with an interferon response. GSEA of DEGs between 3 dpi vs 1 dpi suggested enrichment for the following signaling pathways; “TNFA signaling via NFκB”; “Complement”; “Interferon-gamma”; “Inflammatory response” (**Figure 3I**).

To confirm the secretion of interferon and inflammatory mediators from iPSC-airway infected with SARS-CoV-2, we performed Luminex assays using the basolateral supernatants and apical washes of mock-infected and SARS-CoV-2 infected iPSC-airway **(Figure 4A)**. Consistent with the transcriptomic analysis, IFNβ secretion was increased modestly at 1 dpi in both the apical and basolateral chambers, and increased more drastically by 3 dpi in both chambers. The time-dependent increases in *IL-6, CXCL-9, CXCL-10,* and *TNFα* transcripts were also validated by levels of secreted protein in basolateral media and apical washes **(Figure 4A)**. Though the lack of significant upregulation of CCL2 from 1 to 3 dpi was also confirmed by Luminex assay, there was significant upregulation of GM-CSF in both apical and basolateral chambers. In addition, we observed an increase in TRAIL secretion, suggesting the initiation of apoptosis **(Figure 4A)**. We used RT-qPCR to further validate the key RNA-seq transcriptomic changes in *IFNB1, IFNL1, IFNL2, IFIT1, MX1*, and *IL-6*, confirming an initial modest but significant increase in expression in infected iPSC-airway compared to mock at 1 dpi and a more robust increase by 3dpi **(Figure 4B)**. Of note, the induction of interferon signaling was considerably lower in SARS-CoV-2 infected iPSC-airway compared to Poly(I:C) or recombinant IFNβ treated iPSC-airway **(Figure S3F)**. This result is in line with multiple reports showing that SARS-CoV-2 blocks innate immune signaling^49, 50^. In contrast to iPSC-airway cells, SARS-CoV-2 infected iPSC-derived alveolar type 2 cells (iAT2s) exhibited a delayed and dampened interferon response in our previously published studies^38^.

**Figure 4.**
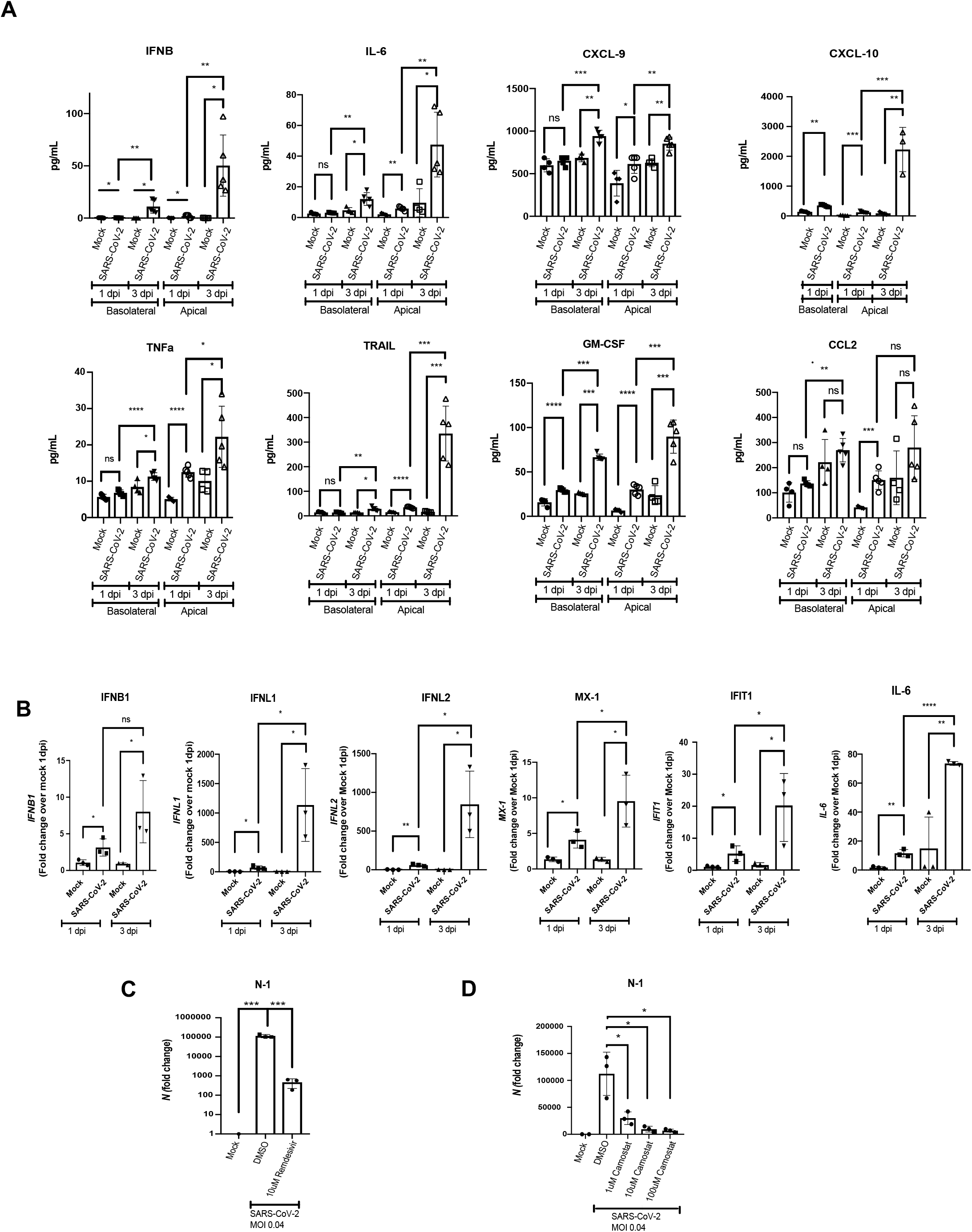
iPSC-airway infected with SARS-CoV-2 secrete inflammatory cytokines and chemokines and detects antiviral drug effects. A) Luminex analysis of apical washes and basolateral media collected from iPSC-airway (BU3 NGPT, cultures (n=3 Transwells). B) RT-qPCR of select genes iPSC-airway (BU3 NGPT) infected with SARS-CoV-2 (MOI 4) and harvested at 1 and 3 dpi with their respective mock-infected samples (n=3 Transwells). Fold change expression compared to mock [2^−ddCt^] after 18S normalization is shown. RT-qPCR of *N* gene expression of mock-infected and SARS-CoV-2 infected (MOI 0.04) iPSC-airway (BU3 NGPT) at 2 dpi pretreated with C) vehicle control (DMSO) or remdesivir (10 μM) or D) vehicle control (DMSO) or camostat (TMPRSS2 inhibitor, 1, 10, 100 μM, as indicated (n= 3 Transwells for both C and D).

We next analyzed the expression of markers of airway epithelial cells, apoptosis, and cell death in our RNA-seq dataset. Markers of multi-ciliated cells (including *FOXJ1*, *TUBA1A* and *DYNLL1*) decreased significantly in the SARS-CoV-2 infected samples by 3 dpi, suggesting loss or perturbation of multi-ciliated cells. Analysis of secretory cell markers demonstrated stable expression of *SCGB1A1* but a relative increase in *MUC5B* at 3 dpi. **(Figure S3E)**. While some markers for cell death were not elevated (BAD), apoptosis markers (*CASP3, PMAIP1, BCL2*) and necrosis/necroptosis markers (*RIPK3*) were upregulated at 3 dpi compared to 1 dpi (**Figure S3E**).

To determine whether the transcriptional response of iPSC-airway was recapitulated in a genetically distinct iPSC line, we performed bulk-sequencing of SARS-CoV-2 infected iPSC-airway at 1 dpi vs mock-infected samples using the 1566 iPSC line. Similar to findings with BU3 NGPT iPSCs, there was evidence of type I and type III interferon response to viral infection by 1 dpi. **(Figure S3C, D)**. GSEA of 1566 infected iPSC-airway revealed almost identical pathway enrichment, including TNF-NFKB, IFN-gamma, IFN-alpha, IL-6/JAK/STAT3 and complement. **(Figure S3B)**.

Taken together, our results indicate that SARS-CoV-2 infected iPSC-airways exhibit transcriptomic changes characterized by significant type I and type III interferon responses, loss of multi-ciliated cell markers, and a pro-inflammatory phenotype characterized by progressively increasing levels of NF-κB signaling. These data are in agreement with other studies in airway and alveolar cultures showing that SARS-CoV-2 infection induces a pro-inflammatory response^14, 38^.

### iPSC-airway can be used as a platform for antiviral drug screening

Finally, to assess the feasibility of iPSC-airway platform to screen for COVID-19 therapeutic compounds, we tested the effect of the FDA-approved antiviral drug remdesivir, and found that viral *N* transcripts were reduced ~100 fold in remdesivir-treated cells **(Figure 4C)**. We also tested the effect of the TMPRSS2 inhibitor camostat mesylate, which significantly reduced the amount of viral N transcripts in a dose-dependent manner (**Figure 4D**). This indicates that SARS-CoV-2 infection of iPSC-airway relies on priming by the protease TMPRSS2, similar to infection in iAT2 cells^38^.

## DISCUSSION

We show here that human iPSC-airway cells derived from multiple individuals express viral entry factors *ACE2* and *TMPRSS2*, are permissive to SARS-CoV-2 infection, and upon infection generate an interferon and inflammatory response. Furthermore, treatment with remdesivir or camostat methylate cause a decrease in viral replication, demonstrating the feasibility of using this iPSC-airway platform for antiviral drug screening assays. Our transcriptomic analyses extend on the findings of a previously published iPSC-derived airway model which was infectable with SARS-CoV-2 and which also showed decrease in infection from 24 to 48 hours, as well as induction of an interferon response^51^. To our knowledge, this is the most detailed assessment of the expression of profile of SARS-CoV-2 entry factors and epithelial response to SARS-CoV-2 infection using iPSC-derived airway cells.

The global burden of respiratory viruses is significant, suggesting the need for in vitro platforms that accurately recapitulate the biology of the human lung. The application of iPSC technology to study viral infections is expanding^52^, with a recent focus on organs infected by SARS-CoV-2^53^. We previously described the application of iAT2s to study infection of the distal lung epithelium by SARS-CoV-2 using^38^. Interestingly, iAT2s were highly permissive to infection, with 20% of cells infected (MOI 5) at 1 dpi and 60% at 4 dpi^38^. Despite similar culture and infection conditions, iPSC-airway cells were less permissive to infection with ~6-12% infected cells at 1 dpi (MOI of 4), and a decrease in viral transcripts by 3 dpi. Consistent with prior studies in primary cells, we detected peak virus release from iPSC-airway, preferentially from the apical surface, at 3 dpi^13, 44^. However, by 3 dpi viral transcripts were decreasing within the epithelium, suggesting either a restriction of viral replication, death of infected cells, or both.

Our transcriptomic analysis revealed that decrease in SARS-CoV-2 transcripts in iPSC-airways between 1 and 3 dpi was accompanied by a significant and progressive induction of both type I and III interferon responses, raising the question of whether the antiviral response in part accounts for the restriction of epithelial infection in our system. Though these results mirror a recent scRNA-seq analysis of SARS-CoV-2 infected primary airways^45^, they contrast with other studies of SARS-CoV-2 infected primary airway cultures that showed minimal type I and type III interferon response^14, 54^. These differences may be due to the observation that iPSC-derived tissues tend to be less mature than primary comparators; whether the robust anti-viral response observed here reflects the biology of young/fetal epithelial tissue will require further study. The robust interferon response in iPSC-airway also differs from the delayed and dampened interferon response of SARS-CoV-2 infected iAT2 cells^38^. These observations give rise to the intriguing speculation that airway epithelial cells may respond differently to SARS-CoV-2 infection than alveolar epithelial cells. In SARS-CoV-1 and MERS-CoV infections, a delayed IFN response in infected human airway epithelial cells was a determinant of disease severity^55, 56 57^. Does the late onset of ARDS in a subset of individuals infected with SARS-CoV-2 develop as a consequence of a failure of the airways to restrict the infection? Future studies aimed at interrogating differential responses of airway and alveolar lung epithelia to SARS-CoV-2 infection may help to address these questions.

Given that hyper-inflammation is a hallmark of SARS-CoV-2 infection and is associated with morbidity/mortality, we also focused on pathways associated with pro-inflammatory cytokine/chemokine production in SARS-CoV-2 infected iPSC-airway. Indeed, SARS-CoV-2 infection was associated with activation of the NFκB pathway as well as chemokines implicated in the recruitment of downstream immune mediators and leukocytes. Specifically, there was a significant upregulation and secretion of IL-6 and TNFα, and T/NK-cell chemokines CXCL9 and CXCL10. There was also an increase in neutrophil-associated chemokines CXCL2 and CXCL8, consistent with the observation that neutrophils have a role in the biology of SARS-CoV-2 infection^58, 59^. IL23, part of TH17 response axis important for mucosal immunity, was modestly upregulated. Though we did not detect a time-dependent increase in monocyte-associated chemokines CCL2 and CCL8 as has been reported in primary airways^45, 54^, we detected an increase in GM-CSF expression from 1 to 3 dpi, suggesting monocyte-macrophage activation may result from epithelial secretion of these cytokines. Our results suggest that iPSC-airway epithelium, upon SARS-CoV-2 infection, is primed to orchestrate downstream immune responses.

In agreement with previous reports on the tropism of SARS-CoV-2 in airway epithelial cells^3, 45, 60^ and in keeping with the tropisms of coronaviruses HCoV-NL63 and SARS-CoV-1^61–63^, we find that multi-ciliated cells are the primary initial airway target cells of SARS-CoV-2. Given lack of convincing co-localization of N+ cells with MUC5AC and TP63, we suspect the majority of N+ cells represent multiciliated cells that had apical acetylated-tubulin on a separate plane as N+ staining. Consistent with this possibility, we observed a time-dependent downregulation of multi-ciliated cell markers during SARS-CoV-2 infection which likewise mirrors observations made in primary cell infection^60^. Significant injury to multi-ciliated cells in this context could impair muco-ciliary clearance, promoting viral spread to the distal airway as well as promoting secondary bacterial infection, both of which are associated with worse outcomes for Covid-19 patients^60, 64^. Primary cell studies have shown that goblet cells eventually become infected with SARS-CoV-2, whereas basal cells are not permissive to infection, possibly due to inaccessibility to the virus due to their basolateral location, and minimal expression of TMPRSS2^2^. We similarly find that iPSC-airway basal cells have minimal expression of TMPRSS2 and ACE2 and are not detectably infected in our model. In individuals with more severe or prolonged infection with SARS-CoV-2 it will be important to determine if basal cells are ultimately infected and understand potential consequences on airway regeneration.

Lastly, we found that treatment with remdesivir or camostat methylate led to a decrease of viral N transcripts in our platform. There remains an urgent need for effective therapies targeting pulmonary viral infections. The iPSC-airway offers an additional physiologically-relevant drug-validating platform to accelerate their development.

This study is not without limitations. The iPSC-airway epithelium is transcriptionally similar to primary airway epithelium and can be used to functionally model diseases such as cystic fibrosis (CF) in vitro^40^. However, iPSC-derived cells differentiated into many cell types of different organs tend to be more fetal or immature than their endogenous counterparts in adults^65–67^, and there may be differences including receptor expression and cell type distributions. Whether our model reflects a more fetal or pediatric response to SARS-CoV-2 will require further investigation. Another limitation of this model is that the epithelial response is studied in isolation, and the complex interplay between epithelial, immune, interstitial, and endothelial cells that leads to COVID-19 pneumonia is not captured in our model. However, the iPSC system offers a reductionist, physiologically relevant model system to study the intrinsic epithelial response and provides key insights into the initial stages of COVID-19. Furthermore, given the clinical spectrum of disease severity caused by SARS-CoV-2 infection there is pressing need to further understand the mechanisms that lead to serve disease. Numerous genes, variants and pathways are implicated in modulating the response to infection and require further investigation^15, 16^. This iPSC-based platform, coupled with gene-editing technology, opens up future directions to evaluate the mechanisms of the airway response to SARS-CoV-2 response.

## EXPERIMENTAL MODEL AND SUBJECT DETAILS

### Maintenance of human iPSCs

All experiments involving the differentiation of human pluripotent stem cell (PSC) lines were performed with the approval of the Institutional Review Board of Boston University (protocol H33122). The two iPSC lines, i) BU3 NGPT, iPSC line carrying NKX2-1-GFP and P63-tdTomato reporter and ii) 1566, a CFTR-corrected line of a homozygous F508del CF iPSC were previously described^40^. BU3 NGPT was derived from the published single reporter, NKX2-1 GFP, iPSC line (BU3 NG), a normal donor iPSC carrying homozygous NKX2-1GFP reporters^21^. The BU3 NG line was then targeted and integrated with a P63tdTomato fluorescent reporter using CRISPR/Cas9 technology. The homozygous F508del iPSC was generated from peripheral blood mononuclear cells at Boston Children’s Hospital Stem Cell Program, and monoallelic correction was performed by nucleofecting the cells with Cas9-GFP plasmid and plasmid containing WT CFTR that includes the exon encoding F508. Repaired clone 1566 (CFTR WT/F508del) was identified by ddPCR and Sanger sequencing. All iPSC lines used in this study displayed a normal karyotype when analyzed by G-banding (Cell Line Genetics). All iPSC lines were maintained in feeder-free conditions, on growth factor reduced Matrigel (Corning) in 6-well tissue culture dishes (Corning), in StemFlex (Gibco), or mTeSR1 medium (StemCell Technologies) using gentle cell dissociation reagent or ReLeSR™(StemCell Technologies) for passaging.

## METHOD DETAILS

### Directed differentiation of human into iPSCs airway epithelium via iBCs

The protocol involves the step-wise directed differentiation of iPSCs to airway as published previously^27, 40^. In brief, it recapitulates the major stages of embryonic lung development as follows: ***Stage 1) Definitive endoderm** (D0-3)* using STEMdiff™ Definitive Endoderm Kit; ***Stage 2) Anterior foregut endoderm** (D3-6)* through TGFβ and BMP inhibition; ***Stage 3) Lung specification** (D6-15)* using a combination of CHIR 99021 (to activate WNT signaling), BMP4, and retinoid acid and evidenced by the expression of NKX2-1. We then purify these NKX2-1+ lung epithelial progenitors at D15 through either sorting on GFP (BU3 NGPT) or for non-reporter line, utilizing a CD47^hi^/CD26^neg^ sorting strategy^21^; ***Stage 4) Early airway organoids** (D15-30),* where GFP+ (BU3 NGPT) or CD47^hi^/CD26^neg^ sorted cells (1566) are plated in 3D Matrigel and cultured in media containing FGF2 and FGF10 for patterning toward proximal airway-like organoids composed of immature secretory and basal progenitors^21, 28^. ***Stage 5) iBCs** (D30-45),* during which the iPSC-3D organoids are further expanded in dual-SMAD inhibition media with DMH-1 (inhibits BMP4-pSMAD 1/5/8) and A83-01 (inhibits TGFβ-pSMAD 2/3) for ~12 days ^68, 69^. iBCs were either serially passaged or cryopreserved, thawed and then serially passaged to generate cells for stage 6. ***Stage 6) Airway differentiation on ALI culture** (>D45)* iBCs are identified and purified by sorting on NGFR+ cells, and purified cells are expanded and differentiated on Transwells in dual-SMAD inhibition media then transitioned to ALI media when >80% confluent. Apical chamber media is removed to initiate ALI to recapitulate a physiologically-relevant environment and stimulate differentiation. After 14 days, iBCs form a pseudostratified airway epithelium that displays morphologic, molecular, and functional similarities to primary human airway epithelial cells and is comprised of the major airway cell types of multi-ciliated, secretory, and basal cells^40^.

### Reanalysis of previously published single-cell RNA-seq dataset for viral entry factors

Viral entry factors *ACE2* and *TMPRSS2* were assessed on scRNA-seq data of 1) BU3 NGPT and HBECs that were previously published^40^ and 2) freshly isolated, uncultured lung epithelia^42^.

For each of the three single cell datasets, previous analyses had performed dimensionality reduction using PCA and UMAP and clusters had been assigned using the Louvain algorithm. We annotated the clusters for each single cell dataset using known markers of secretory, ciliated and basal cells. Following this annotation, the three datasets were merged. When the datasets were merged, in addition to regressing out mitochondrial content during our normalization procedure (SCTransform)^70^, we also regressed out the “library” batch effect. We proceeded to compare the different cell types between the datasets using genes of interest (*ACE2*, *TMPRSS2*, etc) and violin plots. Markers for each dataset were computed using MAST^71^ and direct comparisons were made between each of the datasets in order to quantitatively assess the differences in expression between key genes (*ACE2*, *TMPRSS2*, etc). All single cell visualizations were made with Seurat (heatmaps, UMAPs, violin plots)^72, 73^. Positive cells were identified using a UMI threshold of 0 counts, so that any cell with 1 UMI or more for a specific gene was considered positive for that gene.

### SARS-CoV-2 propagation and titration

SARS-CoV-2 stocks (isolate USA_WA1/2020, kindly provided by CDC’s Principal Investigator Natalie Thornburg and the World Reference Center for Emerging Viruses and Arboviruses (WRCEVA)) were grown in Vero E6 cells (ATCC CRL-1586) cultured in Dulbecco’s modified Eagle’s medium (DMEM) supplemented with 2% fetal calf serum (FCS), penicillin (50 U/mL), and streptomycin (50 mg/mL) and titrated as described previously^38^. To remove confounding cytokines and other factors, viral stocks were purified by ultracentrifugation through a 20% sucrose cushion at 80,000xg for 2 hours at 4°C as previously published^38^. SARS-CoV-2 titer was determined in Vero E6 cells by tissue culture infectious dose 50 (TCID50) assay. All work with SARS-CoV-2 was performed in the biosafety level 4 (BSL-4) facility of the National Emerging Infectious Diseases Laboratories at Boston University following approved SOPs.

### SARS-CoV-2 infection of iPSC-airway on ALI culture

Prior to infection, the apical side of iPSC-airway plated in ALI culture was washed with 100 μL of PBS for 5-10 minutes at 37°C to remove the mucus. Then, purified SARS-CoV-2 stock (100 μL of inoculum prepared in PBS) or PBS without virus (mock infection) was added at the apical chambers of each Transwell at the indicated multiplicity of infection (MOI) and allowed to adsorb for 1 hour at 37°C and 5% CO_2_. After adsorption, the inoculum was removed and cells were incubated at ALI at 37°C for 1-7 days. Basolateral chamber medium was changed every 2-3 days. To ensure that the levels of *N*-transcript were due to infected epithelial cells and not virions on the surface of the epithelium, we compared levels of *N* transcript at 1 dpi to day 0 cultures which had been exposed to SARS-CoV-2 for ~10 minutes, and saw more *N* in 1 dpi samples for both BU3 NGPT and 1566 (**Figure S2B**). At the time of harvest, basolateral media was collected for further analysis. Apical washes were performed by adding 100 μL media to the apical chamber and incubation for 15 minutes at 37°C before collection for further analysis. Both the apical washes and basolateral media were used for viral titration and Luminex assays as described below. For immunofluorescent analysis or electron microscopy, cells were fixed in 10% formalin. For RT-qPCR and RNA-seq analysis, cells were harvested directly in TRIzol (ThermoFisher).

### Cell viability assay

For determining cell viability, iPSC-airways cultured at ALI were detached by adding 0.2 mL Accutase apically and 0.5 mL basolaterally and incubated at 37°C for 15 minutes. Detached cells were pelleted by low-speed centrifugation, resuspended in PBS, diluted 1:1 in trypan blue, and analyzed using a LUNA-II™ Automated Cell Counter (Logos Biosystems).

### Immunofluorescence microscopy of iPSC-airway

SARS-CoV-2 infected or mock-infected cultures on Transwell inserts were fixed in 10% formalin for 6 hours, washed twice in PBS (10 minutes each, room temperature), permeabilized with PBS containing 0.25% Triton X-100 and 2.5% normal donkey serum (30 minutes, room temperature), and blocked with PBS containing 2.5% normal donkey serum (20 minutes, room temperature). Subsequently, cells were incubated with primary antibody diluted in 4% normal donkey serum overnight at 4°C. The antibodies used were; anti-SARS-CoV nucleoprotein (N) antibody (rabbit polyclonal, 1:2500, Rockland Immunochemicals, Cat #200-401-A50), anti-α-TUBULIN antibody (mouse monoclonal, 1:500 sigma cat# T6199), and anti-MUC5B antibody (mouse monoclonal, Santa Cruz Biotechnology, 1:500 cat#SC-39395-2). The anti-SARS-CoV nucleoprotein (N) antibody cross-reacts with the SARS-CoV-2 nucleoprotein^38^. Next, cells were washed with PBS three times (5 minutes, room temperature), and incubated with secondary antibody (AlexaFluor 546 AffiniPure Donkey Anti-mouse IgG (H+L), 1:500, and AlexFluor 647 donkey anti-mouse IgG (H+L) 1:500, Jackson ImmunoResearch) for 2 hours at room temperature. Cells were washed with PBS three times (5 minutes, room temperature), incubated with DAPI (1:5000, Life Technologies) for 5 minutes, and washed again. Transwell inserts were then cut out with a razorblade and mounted with Prolong Diamond Mounting Reagent (Life Technologies). Slides were imaged with either upright fluorescence microscope (Nikon Eclipse Ni-E) or confocal microscope (Leica SP5). Manual quantification of N+ cells were performed by distributing 10-15 images of iPSC-airway infected with SARS-CoV-2 stained with DAPI and N (>200 cells/image) to 5 blinded scorers. Each image analyzed using ImageJ and DAPI+ nuclei and N+ cells were quantified with the multi-point tool. To determine cellular tropism, Z-stack images of the infected transwells stained with either anti-SARS-CoV-2 N and α antibodies or anti-SARS-CoV-2 N and anti-mucin 5B antibodies were taken on Nikon Eclipse NiE. For quantification of co-localization of N+ cells with airway cell types, at least 100 N+ cells were counted, and percentage of co-localization with α-tubulin and mucin 5B were determined.

### ACE2 Immunohistochemistry

For ACE2 immunohistochemistry, iPSC-airways on Transwells were fixed in 4% PFA for 30 minutes, washed three times with PBS (5 minutes each, room temperature), then dehydrated in 50% ethanol (15 minutes), 70% ethanol (15 minutes), 85% ethanol (15 minutes), 95% ethanol (15 minutes), 100% ethanol (three times, 15 minutes each), and Histoclear (Biocare Medical, 3 times 15 minutes each), and incubated in paraffin (3 changes 30 minutes each) at 60°C. The filter were cut out of the Transwells, then cut in half, and embedded in paraffin. The embedded filters were sectioned on a Leica RM 2135 microtome at a thickness of 7 μM. Sections were deparaffinized, blocked with Inhibitor CM and antigen retrieval was performed with Cell Conditioner 1 (CC1). Anti-ACE2 (Abcam cat# ab108252) was applied and was visualized with anti-rabbit HQ and anti-HQ-HRP followed by ChromoMap DAB (Roche). Samples were counter stained with hematoxylin, rinsed with detergent, dehydrated, and coverslipped with permanent mounting media. For MUC5AC co-staining, prior to the incubation of the second primary ab, the samples were subjected to a heat denature step with Cell Conditioner 2 (CC2). Anti-MUC5AC (Abcam cat# ab198254) was applied and visualized with anti-rabbit NP (Roche) and anti-NP-AP (Roche) and detected with Discovery Yellow (Roche); counter stained with hematoxylin, rinsed with detergent, dehydrated, and cover-slipped with permanent mounting media. For tubulin co-staining, sections were deparaffinized, blocked with Inhibitor CM and antigen retrieval was performed with CC1. Anti-ACE2 (Abcam cat# ab108252) was applied and was visualized with anti-rabbit HQ and anti-HQ-HRP followed by ChromoMap DAB (Roche). Prior to the incubation of the second primary ab, the samples were subjected to a heat denature step with CC2. Anti-alpha Tubulin (Abcam cat# ab24610) was applied and visualized with anti-mouse OmniMap-HRP (Roche) and detected with Discovery Purple (Roche) counter stained with hematoxylin, rinsed with detergent, dehydrated, and a cover-slip placed with permanent mounting media.

### Reverse Transcriptase Quantitative PCR (RT-qPCR)

RNA was extracted from TRIzol samples following the manufacturer’s protocol. Purified RNA was reverse transcribed into cDNA using the MultiScribe Reverse Transcriptase (Applied Biosystems). All qPCR was performed in 384-well plates and run for 40 cycles using an Applied Biosystems QuantStudio 384-well system. Predesigned TaqMan probes were from Applied Biosystems or IDT. Relative gene expression was calculated based on the average Ct value for technical triplicates, normalized to 18S control, and fold change over mock-infected cells was calculated using 2^−ΔΔCt^. If probes were undetected, they were assigned a Ct value of 40 to allow for fold change calculations, and replicates, as indicated in each figure legend, were run for statistical analyses. A replicate of the RT-qPCR is defined as an individual Transwell of airway epithelial cells generated from sorted NGFR+ cells from stage 6 of the differentiation protocol.

### Transmission electron microscopy

iPSC-airway (1566) on Transwell inserts were infected with SARS-CoV-2 at an MOI of 4 or mock-infected. At 1 dpi, cells were fixed in Karnovsky’s fixative (Tousimis) for 18 hours at 4°C and washed with PBS. The membrane was excised from the Transwell, block stained in 1.5% uranyl acetate (Electron Microscopy Sciences, EMS) for 1 hour at room temperature (RT). The samples were dehydrated quickly through acetone on ice, from 70% to 80% to 90%. The samples were then incubated 2 times in 100% acetone at RT for 10 minutes each, and in propylene oxide at RT for 15 minutes each. Finally, the samples were changed into EMbed 812 (EMS), left for 2 hours at RT, changed into fresh EMbed 812 and left overnight at RT, after which they were embedded in fresh EMbed 812 and polymerized overnight at 60°C. Embedded samples were thin sectioned (70 nm) and grids were stained in 4% aqueous uranyl acetate for 10 minutes at RT followed by lead citrate for 10 minutes at RT. Electron microscopy was performed on a Philips CM12 EM operated at 100kV, and images were recorded on a TVIPS F216 CMOS camera with a pixel size of 0.85-3.80 nm per pixel.

### Drug efficacy testing in iPSC-airway cells

iPSC-airway plated in ALI culture were pre-treated apically (100 μL) and basolaterally (600 μL) with the indicated concentrations of camostat mesylate (Tocris, #59721-29-5), remdesivir (Selleckchem, #S8932), or DMSO control for 1 hour at 37°C. After 1 hour, all apical media were aspirated and SARS-CoV-2 (MOI 0.04) was added for 1 hour without any drugs apically, after which the inoculum was removed, and samples were returned to 37°C in ALI culture format. iPSC-airway were exposed to the compounds basolaterally for the entire duration of the experiment. Cells were harvested in TRIzol 2 dpi and processed for RT-qPCR.

### Immune stimulation with poly(I:C) and IFNβ

For immune stimulation treatments, iPSC-airways cultured at ALI were treated with the TLR3 agonist poly(I:C) (InvivoGen) delivered with Oligofectamine Transfection Reagent (Invitrogen) or treated with recombinant human IFNβ (rhIFNβ) (PeproTech). Prior to treatment, 1 μL poly(I:C) was mixed with 2.5 μL Oligofectamine and incubated at RT for 15 minutes. After the incubation period, the poly(I:C) and Oligofectamine mixture was added to 100 μL of ALI differentiation media. iPSC-airway cultured at ALI were treated apically (100 μL) with poly(I:C) (10 μg/mL) and Oligofectamine, or apically and basolaterally (600 μL) with IFNβ (10 ng/mL) for 24 hours at 37°C. Cells were subsequently harvested in TRIzol for RT-qPCR.

### Luminex analysis

Apical washes and basolateral media samples were clarified by centrifugation and analyzed using the Magnetic Luminex^®^ Human Discovery Assay (R&D Systems). Custom configured targets include: CCL2/JE/MCP-1, CXCL-9/MIG, CXCL10/IP-10/CRG-2, GM-CSF, IFNβ, IL-6, TNFα, TRAIL/TNFSF10. Mean fluorescence intensity was measured to calculate final concentration in pg/mL using Bioplex200 and Bioplex Manager 5 software (Biorad).

### RNA sequencing and bioinformatic analyses

For bulk RNA sequencing (RNA-Seq), biological triplicate (n=3) samples of purified RNA extracts were harvested from each group of samples prepared as follows. After 87 days of total time in the directed differentiation protocol, iPSC-airways cultured as serially passaged 3D spheres were sorted and single-cell passaged onto Transwell inserts. Apical media was removed on day 8 to initiate ALI culture. On day 24 (after removing the apical media), 6 replicate wells of iPSC-airways were infected with SARS-CoV-2 from the apical surface and 6 replicate wells were mock-infected. Three wells per condition were harvested at each 1 and 3 dpi in TRIzol. Following total RNA isolation from the TRIzol samples, mRNA was isolated from each sample using magnetic bead-based poly(A) selection, followed by cDNA synthesis. The products were end-paired and PCR-amplified to create each final cDNA library. Sequencing of pooled libraries was done using a NextSeq 500 (Illumina). The quality of the raw data was assessed using FastQC v.0.11.7^74^.The sequence reads were aligned to a combination of the human genome reference (GRCh38) and the SARS-CoV-2 reference (NC_045512) using STAR v.2.6.0c ^75^. Counts per gene were summarized using the featureCounts function from the subread package v.1.6.2^76^. The edgeR package v.3.25.10^77^ was used to import, organize, filter and normalize the counts. The matrix of log counts per million was then analyzed using the limma/voom normalization method^78^. Genes were filtered based on the standard edgeR filtration method using the default parameters for the “filterByExpr” function. After exploratory data analysis with Principal Component Analysis (PCA), contrasts for differential expression testing were done for each SARS-CoV-2-infected sample vs mock (controls) at each time point (days post infection). Differential expression testing was also conducted to compare the gene expression between the two infected time points and to investigate the time specific effects in response to infection. The limma package v.3.42.2^78^ with its voom method, namely, linear modelling and empirical Bayes moderation was used to test differential expression (moderate t-test). P-values were adjusted for multiple testing using Benjamini-Hochberg correction (false discovery rate-adjusted p-value; FDR). Differentially expressed genes for each comparison were visualized using Glimma v. 1.11.1^79^ and FDR<0.05 was set as the threshold for determining significant differential gene expression. Functional predictions were performed using the fgsea v.1.12.0 package for gene set analysis^80^.

## QUANTIFICATION AND STATISTICAL ANALYSIS

Statistical analyses were performed using GraphPad Prism 8, ggplot2 R package. Statistical significance was determined as P-value of < 0.05 using Student’s t test.

## DATA AND CODE AVAILABILITY

The RNA-Seq data discussed in this publication have been deposited in NCBI’s Gene Expression Omnibus (GEO) and are accessible through GEO Series accession number GSE153277 (reviewer token sjqdimeipxyjtqz).

## Acknowledgements

We thank Brian R. Tilton of the BUSM Flow Cytometry Core and Yuriy Alekseyev of the Boston University School of Medicine (BUSM) Sequencing Core; supported by NIH grant 1UL1TR001430. For facilities management, we are indebted to Greg Miller (CReM Laboratory Manager) and Marianne James (CReM iPSC Core Manager) supported by grants N01 75N92020C00005 and U01TR001810. The current work was supported by Harry Shwachman Cystic Fibrosis Clinical Investigator Award, Gilead Sciences Research Scholars, Alfred and Gilda Slifka Fund, and CF/MS fund to RW; JL was supported by T32 HL007035; F30HL147426 to KMA. R01HL122442, R01HL095993, U01HL134745, U01HL134766 to DNK, U01HL148692 to DNK and FJH; Evergrande MassCPR awards to DNK and EM. R01HL139799 to FJH. iPSC distribution and disease modeling is supported by NIH grants U01TR001810 and NO1 75N92020C00005.

## Author contributions

RW, AH, EM, DK and FH conceived the work, designed the experiments and wrote the manuscript. RW, MLB, CSR, JL, JH, RW and KMA performed the directed differentiation experiments and with AH and EB developed the SARS-CoV-2 infection strategy. AH performed the SARS-CoV-2 infections. JO performed the Luminex analysis. JLV and CVM analyzed the scRNA-Seq and RNA-Seq datasets. MG provided ACE2 immunohistochemistry. AAW provided critical input. AH prepared the samples for TEM and EB performed the TEM and provided analysis support.

---

**Figure S1.**
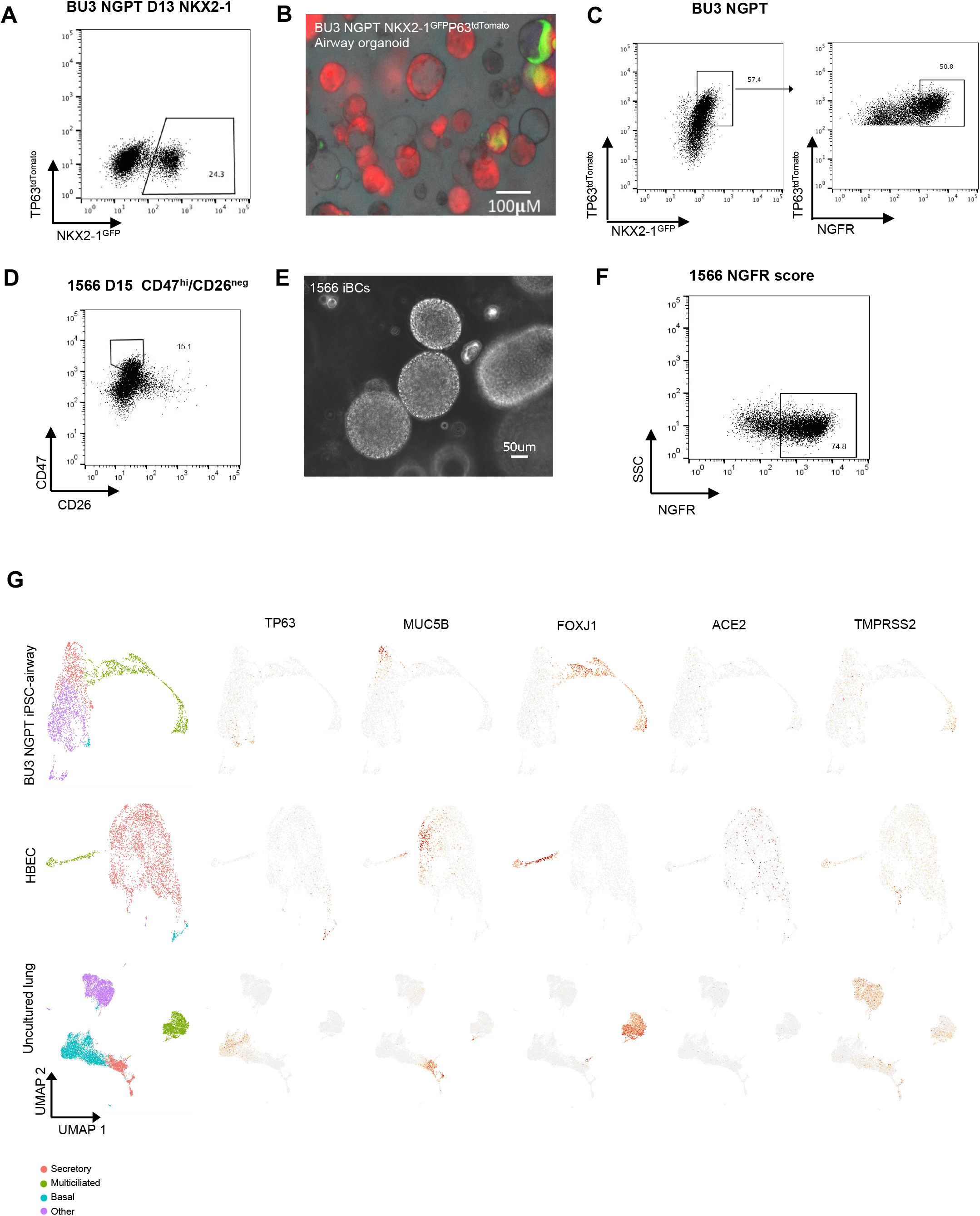
(related to Figure 1). Directed differentiation of human iPSCs to airways epithelium and scRNA-Seq analysis of iPSC-airways, primary HBECs and uncultured lung epithelia. A) Flow cytometry analysis of Day 15 differentiation of BU3 NGPT using the schematic in 1A. In the example show 24.3% cells are NKX2-1^GFP+^ and no cells co-express NKX2-1^GFP^ and TP63^tdTomato^. B) Representative microscopy images of BU3 NGPT at D28. The image is a merge of phase contrast, tdTomato and GFP fluorescence (scale bar =100 μm). C) Representative flow cytometry analysis of BU3 NGPT iBCs analyzing the expression of NKX2-1^GFP+^, TP63^tdTomato^, and NGFR. D) Flow cytometry analysis of Day 15 differentiation of 1566 using CD47^hi^/CD26^neg^ sorting strategy to purify NKX2-1+ lung progenitor cells. In the example show, 15% of cells are selected as CD47^hi^/CD26^neg^ E) Representative 1566 D40+ iBCs (scale bar = 50 μm). F) Representative FACS of 1566 differentiation showing NGFR+ population. G) UMAP of scRNA-seq data from iPSC-airway (BU3 NGPT)^40^, HBEC^40^, and uncultured lung^42^ after Louvain clustering. Clusters were annotated based on the expression of canonical markers and top differentially expressed genes. Examples of key airway cell-specific markers *TP63*, *MUC5B*, *FOXJ1*, as well as viral entry factors *ACE2* and *TMPRSS2* from each sample are shown.

**Figure S2.**
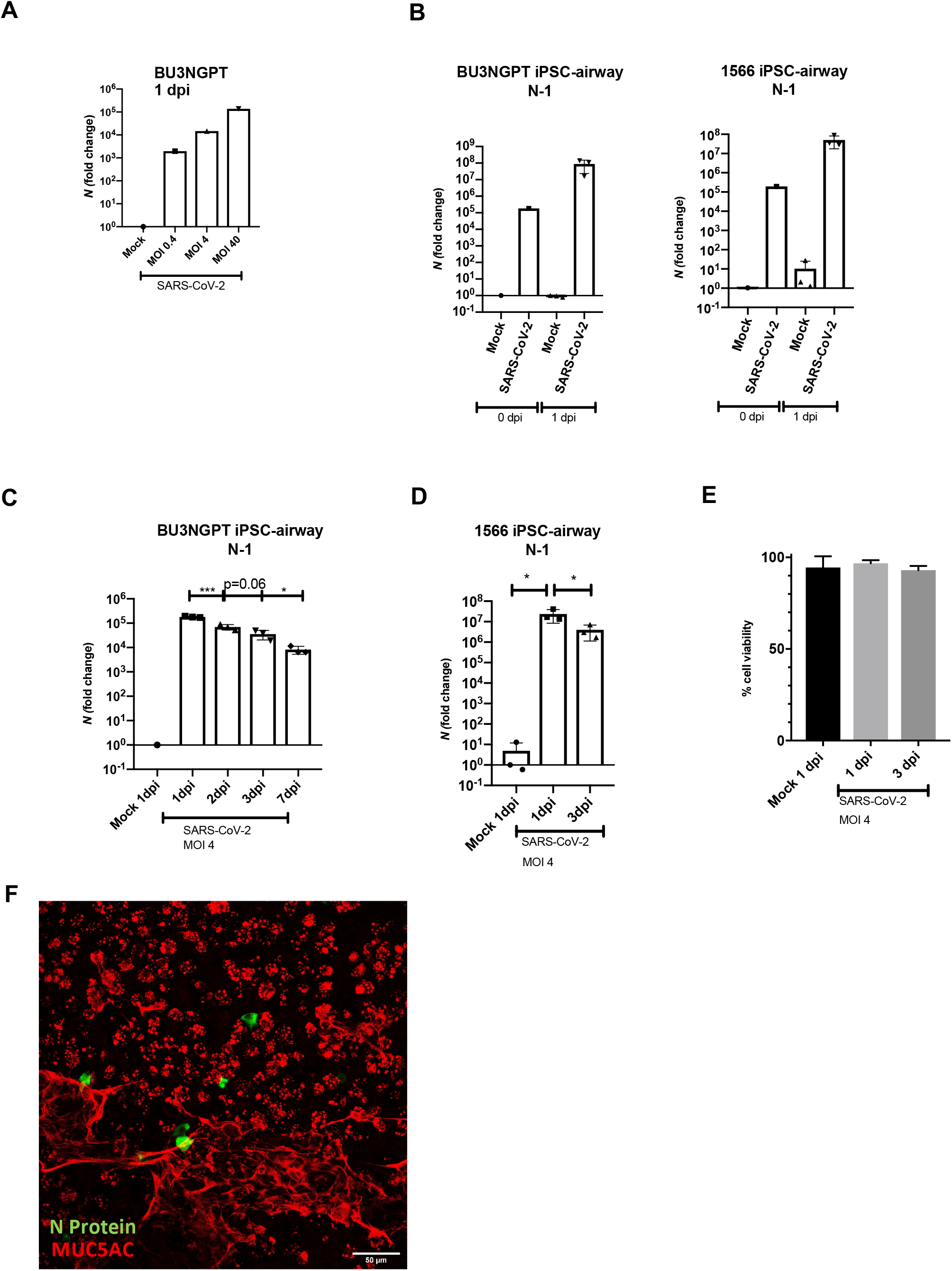
(related to Figure 2). iPSC-airway are permissive to SARS-CoV-2 infection and show time-dependent restriction in viral growth. A) RT-qPCR of viral nucleocapsid *N* gene expression of iPSC-airway (BU3 NGPT) infected with a range of MOIs (0.4, 4, 40) of SARS-CoV-2 analyzed at 1 dpi (n=1 Transwell at each MOI). Fold change expression compared to mock [2^−ddCt^] after 18S normalization is shown. MOI 4 used for all other experiments. B) RT-qPCR of N gene expression of iPSC-airway (BU3 NGPT and 1566) infected with SARS-CoV-2 at 0 (n=1) and 1 dpi (n=3 Transwells). C) RT-qPCR of N gene expression of iPSC-airway (BU3 NGPT) infected with SARS-CoV-2 at 1, 2, 3, and 7 dpi (n=3 Transwells each time point). D) RT-qPCR of N gene expression of iPSC-airway (1566) infected with SARS-CoV-2 from 1 to 3 dpi (n=3 Transwells from each time-point) E) Percentage of viable iPSC-airway (BU3 NGPT) mock infected or infected with SARS-CoV-2 at 1 and 3 dpi as determined by trypan blue staining. F) Confocal microscopy of iPSC-airway (BU3 NGPT) infected with SARS-CoV-2 at 1 dpi stained for SARS-CoV-2 N and MUC5AC (Scale bar = 50μm).

**Figure S3.**
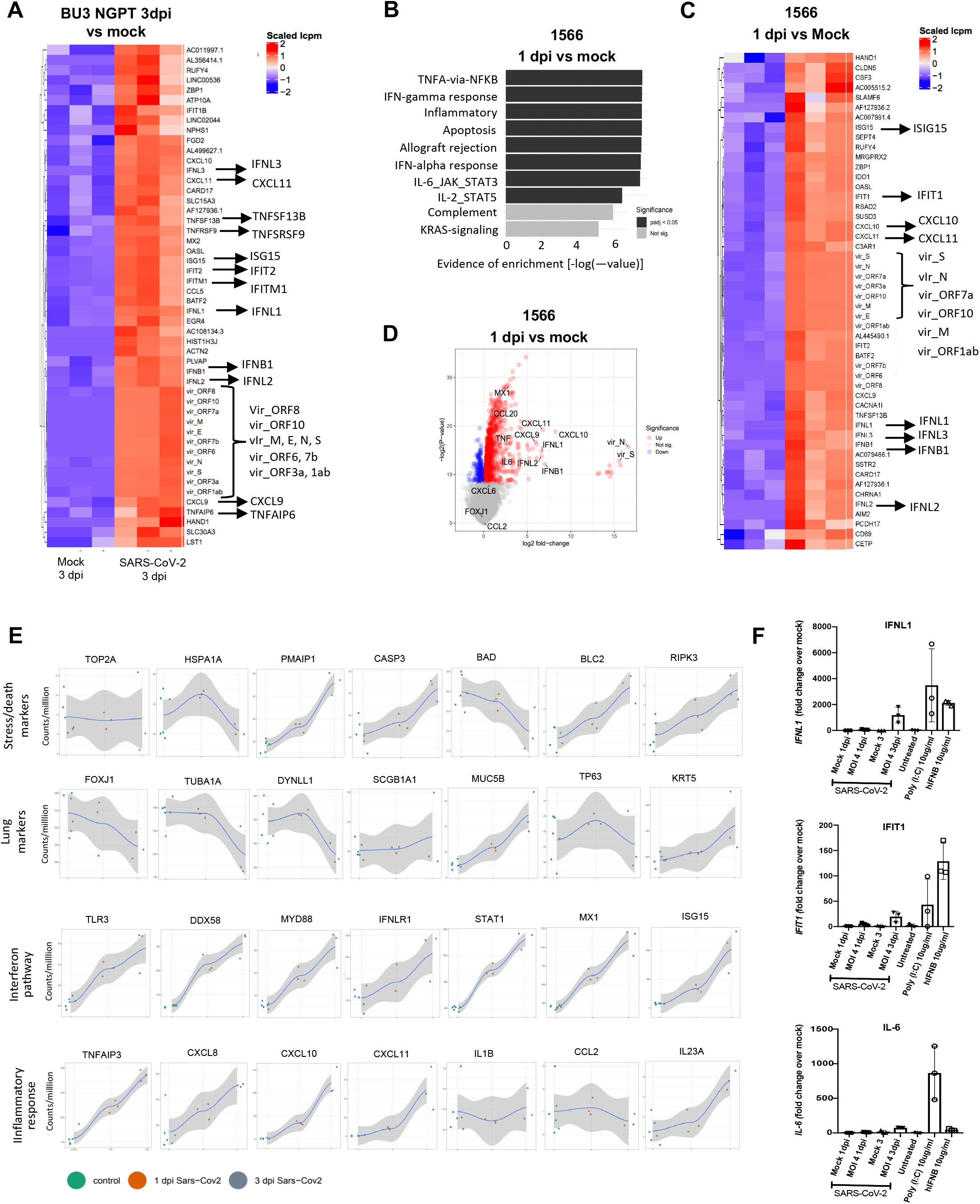
(related to Figure 3). Transcriptomic analysis of SARS-CoV-2 infected iPSC-airway shows a rapid and robust interferon response. A) Unsupervised hierarchical clustered heatmaps of differentially expressed genes (DEGs; −log2FC) between mock-infected and SARS-CoV-2 infected iPSC-airway (BU3 NGPT) samples and samples at 3 dpi. B) Gene set enrichment analysis (GSEA) of the top ten upregulated gene sets in mock versus SARS-CoV-2 (infected iPSC-airway (1566) at 1 dpi. C) Unsupervised hierarchical clustered heatmaps of differentially expressed genes (DEGs; −log2FC) between mock versus SARS-CoV-2 infected iPSC-airway (1566) samples at 1 dpi. D) Volcano plots of differentially expressed genes in mock versus SARS-CoV-2 infected iPSC-airway (1566) at 1 dpi. E) Local regression (LOESS) plots of viral, interferon and ISG, and inflammatory gene expression levels quantified by RNA-seq normalized expression (counts per million reads) for iPSC-airway (BU3 NGPT). F) RT-qPCR of IFNL1, IFIT1, and IL6 in iPSC-airway (BU3 NGPT) that have been mock infected or infected with SARS-CoV-2 (MOI 4) at 1 dpi compared to poly(I:C) transfection (10 μg/mL), or treatment with recombinant human IFNβ(10 ug/mL) for 24 hours. Fold change expression compared to mock [2^−ddCt^] after 18S normalization is shown.

**Table S1.**
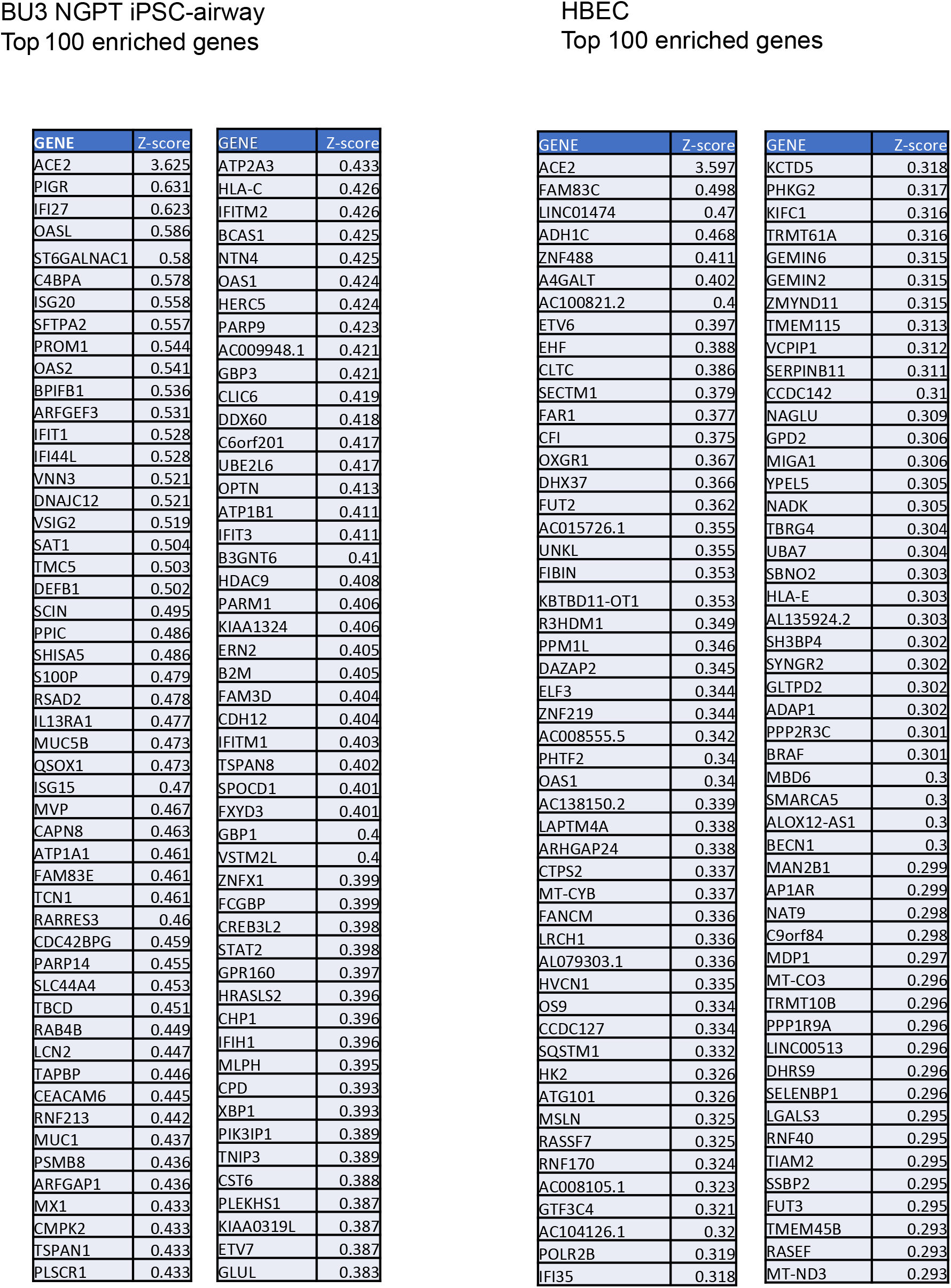
(related to Figure 1). Top 100 enriched genes of ACE2+ cells from sc-RNAseq iPSC-airway (BU3 NGPT) and HBEC^40^ (Hawkins et al, CSC, 2020)

**Table S2. Table of differentially expressed genes (DEGs)in iPSC-airway after SARS-CoV-2 infection**. Listed are the names and statistics (fold-change and false-discovery rate-adjusted values; FDR) for all genes tested through bulk RNA-seq analysis of iPSC-airway infected with SARS-CoV-2 (MOI 4) at 1 and 3 dpi, along with mock-infected cells at 1 and 3 days. Biostatistical comparisons were performed between infected and mock conditions at both 1 dpi and 3 dpi, and the infected time points (3 dpi and 1 dpi), with an expression cut-off for significant differential expression of FDR < 0.05. We have also included normalized expression as log counts per million for each of the samples.

